# Bimodal Spindle Orientation Drives Tissue Regularity in a Proliferating Epithelium

**DOI:** 10.1101/178517

**Authors:** Tara M. Finegan, Daxiang Na, Austin V. Skeeters, Nicole S. Dawney, Patrick W. Oakes, Alexander G. Fletcher, Dan T. Bergstralh

## Abstract

We investigated the relationship between proliferation and tissue topology in an epithelial tissue undergoing elongation. We found that cell division is not required for elongation of the early *Drosophila* follicular epithelium, but does drive the tissue towards optimal geometric packing. To increase tissue regularity, cell divisions are oriented in the planar axis, along the direction of tissue expansion. Planar division orientation is governed by apico-cortical tension, which aligns with tissue expansion but not with interphase cell shape elongation. Hertwig’s Rule, which holds that cell elongation determines division orientation, is therefore broken in this tissue. We tested whether this observation could be explained by anisotropic activity of the conserved Pins/Mud spindle-orienting machinery, which controls division orientation in the apical-basal axis. We found that Pins/Mud does not participate in planar division orientation. Rather, tension translates into planar division orientation in a manner dependent on Canoe/Afadin, which links actomyosin to adherens junctions. These findings demonstrate that division orientation in different axes - apical-basal and planar - is controlled by distinct, independent mechanisms in a proliferating epithelium.

**Summary Statement:** Regularity in a proliferating epithelium requires cells to divorce division orientation from interphase shape, which they accomplish by using distinct mechanisms to orient divisions in the apical-basal and planar axes.

## Introduction

Epithelial morphogenesis is driven by cell-intrinsic and -extrinsic influences that stimulate diverse combinations of cellular behaviours, including rearrangement, shape change, and division. For example, the germband tissue of the early *Drosophila* embryo utilizes both polarized cell rearrangements and cell shape changes to converge (narrow) and extend the tissue initially, with a later contribution from oriented cell division (reviewed in (Lye and Sanson, 2011)). In contrast, expansion of the pupal notum is driven by the combined activity of cell division, rearrangement, shape changes and cell extrusion (Guirao et al., 2015). Whereas development in the early embryo and pupal notum involves complex changes in tissue shape and/or cell patterning, morphogenesis of the early *Drosophila* follicular epithelium (FE) – a simple, cuboidal monolayer - is characterized by gradual and apparently uniform expansion. Because expansion has a directional bias, the FE is a straightforward *in vivo* model for the relationship between elongation and morphogenetic cell behaviour.

The FE surrounds the sixteen germline cells of an egg chamber during oogenesis. As it matures, the egg chamber increases in volume while simultaneously elongating (Figure 1A). Elongation initially relies on the FE, which secretes a basement membrane and then migrates along it by building and breaking adhesions in a planar polarized manner, driving rotation of the egg chamber (Figure 1A’) (Barlan et al., 2017; Cetera et al., 2014; Haigo and Bilder, 2011). The basement membrane is stronger over the elongating bulk of the egg chamber than the poles, and this gradient of stiffness biases expansion in one direction (Crest et al., 2017). The FE is proliferative as early changes in egg chamber shape and size take place. By convention, maturation of the egg chamber is divided into 14 stages. Between developmental stages 2 and 6/7, when mitoses cease, 4-5 rounds of division result in approximately 900 follicle cells (Duhart et al., 2017a; Horne-Badovinac and Bilder, 2005; Klusza and Deng, 2011; Kolahi et al., 2009).

**Figure 1:**
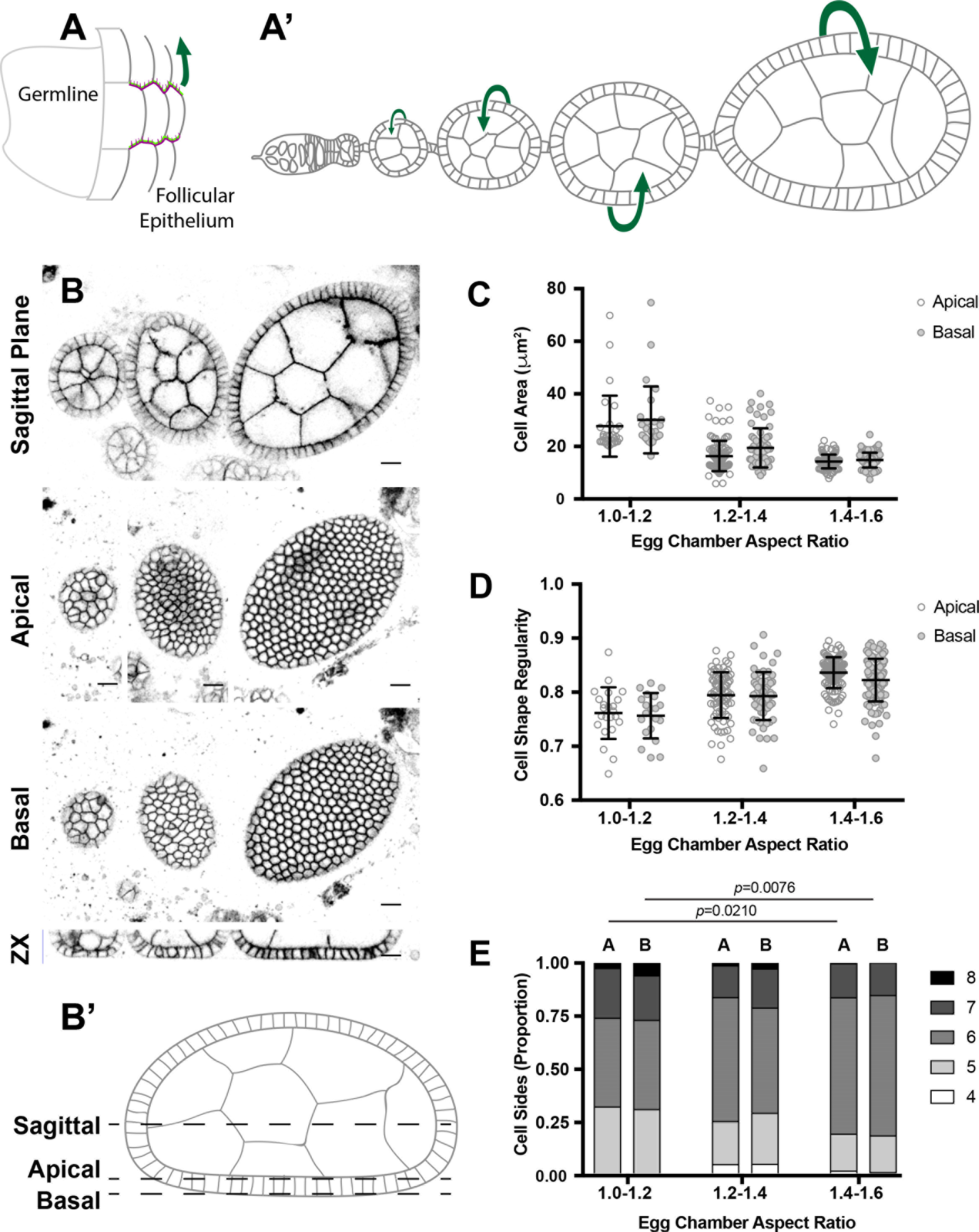
Epithelial topology becomes increasingly regular as the early egg chamber matures. **A)** Collective migration of the follicular epithelium (A’) drives rotation of early egg chambers, which elongate as they increase in size (A). **B)** Egg chambers flatten along the coverslip, allowing for follicle cell morphologies to be compared in two dimensions. In (B), three live *w*^*1118*^ (control) egg chambers expressing the membrane marker Basigin::YFP are shown in multiple planes of focus, with a 3-dimensional z-reconstruction at the bottom. Because the egg chambers flatten at the coverslip, the basal follicle cell morphologies can be revealed in a single focal plane. To account for differences in follicle cell height between the three egg chambers, which increase in maturity from left to right, separate focal planes are shown for the apical morphologies. A diagram illustrating the position of the three focal planes with respect to the flattened egg chamber is shown in (B’). Scale bars = 20μm. **C, D, E)** Follicle cell cross-sectional area decreases (C) and cell shape becomes more regular (D) and hexagonal (E) as early-stage egg chambers mature. These measurements were derived from at least three egg chambers from at least three flies per AR range. Bars in (C) and (D) represent mean and standard deviation. We used cell circularity:

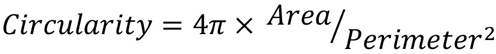

as a measure of cell shape because it depends on both cell perimeter and area, and is therefore more sensitive to deviations than aspect ratio alone. Statistical significance for cell side distributions was calculated using a chi-squared test. Number of cells used for morphological analysis: 1.0-1.2 apical n =23, basal n=27; 1.2-1.4 apical n=114, basal n=61; 1.4-1.6, apical n=160, basal n=149. Vertex counts 1.0-1.2 apical n =38, basal n=33; 1.2-1.4 apical n=73, basal n=70; 1.4-1.6, apical n=284, basal n=259.

These divisions are oriented with respect to the apical-basal axis, as they are in most epithelia, such that new cells are born within tissue layer. Apical-basal division orientation is governed by a highly-conserved cortical machinery that includes Pins (Partner of Inscuteable) and Mud (Mushroom body defective). Mud uses dynein to exert a pulling force on astral microtubules, and thereby draws the mitotic spindle into alignment (reviewed in (Bergstralh et al., 2017; di Pietro et al., 2016)). Division orientation with respect to the plane of the tissue has not been examined in the FE, but is determined in other systems by asymmetric localization of the Pins/Mud machinery (Bosveld et al., 2016; Hart et al., 2017).

In this study, we addressed the question of how epithelial topology develops in an elongating tissue. We determined that the tissue becomes increasingly regular as a consequence of proliferation, and that proliferation relieves stress exerted on the tissue by elongation. To achieve both of these goals, follicle cells in longer egg chambers must break Hertwig’s Rule, orienting divisions contrary to the direction predicted by interphase cell shape (Hertwig, 1884). Follicle cells achieve planar division orientation through a Mud-independent mechanism that relies on the mechanosensing of apical tension.

## Results and Discussion

### The Follicle Epithelium Becomes Increasingly Regular Over Proliferating Stages of Egg Chamber Development

Follicle cell geometry was examined in pre-vitellogenic egg chambers, which have aspect ratios (AR - long axis / short axis) between approximately 1.0 and 1.6 (Cetera et al., 2014). We took advantage of the fact that live egg chambers imaged in oil flatten perceptibly as they settle onto the coverslip (Figure 1B and B’), providing an area in which cells can be compared in two dimensions. This area is restricted to the central bulk of the egg chamber, over which elongation occurs, and excludes the terminal poles.

We used automated segmentation and image analysis to measure features of cell and tissue geometry, and found that tissue topology becomes more regular as the egg chamber elongates (Figures 1D and S1A). At the earliest stages of egg chamber development, follicle cells exhibit a wide range of morphologies (Figure 1C–E). We also frequently observed that cell height was uneven in these chambers, indicative of localized differences in tension (Figure S1B). These observations suggest that the tissue is in mechanical disequilibrium when the egg chamber leaves the germarium.

By approximately stage 6/7 (AR 1.4-1.6), cells are consistently smaller, meaning that proliferation outpaces elongation. Cells are also more likely to be hexagonal (Figure 1C-E). We considered the possibility that cell geometry could change along the apical-basal (A-B) axis of the tissue, and therefore measured near to both the apical and basal tissue surfaces, but found that the parameters we measured did not change significantly between them (Figure 1B). These measurements demonstrate that the follicular epithelium transitions from a loosely organized state in very early egg chambers to a classic well-packed hexagonal ‘honeycomb’ by the time division stops at stage 6/7 (Thompson, 2014).

### Tissue Regularity, but not Elongation, Relies on Cell Division

These observations raise the question of how increased tissue regularity is coordinated with elongation. We first considered the possibility that elongation and regularity are driven by increased junctional tension, a function of contractility resulting from actomyosin and cell-cell adhesion (reviewed in (Lecuit et al., 2011)). We found that programmed modulation of junctional tension is unlikely to drive regularity in the follicle epithelium, since we failed to see changes in the amount of either Armadillo (*Drosophila* β-catenin) or Zipper (*Drosophila* Myosin Heavy Chain II) at the apical surface over the course of early egg chamber elongation (Figures S2A,B). We also noticed distinct spots, or “blobs”, of Zipper positioned along some follicle cell lateral membranes. These blobs are reminiscent of somatic cell ring canals, which arise from incomplete cytokinesis and might therefore be expected to demonstrate enrichment of Myosin. Consistent with this, we observed that Squash (*Drosophila* Myosin Regulatory Light Chain II) colocalizes with the ring canal marker Pavarotti along lateral membranes (Figure S2C) (Airoldi et al., 2011). We did not observe any pattern suggesting an active role for ring canals in epithelial topology.

These observations suggest that elongation and regularity in the FE are passive processes. We therefore addressed the importance of proliferation to topology. The finding that proliferation outpaces directional growth of the egg chamber (Figure 1C) led us to ask whether extra cells are needed to drive egg chamber rotation, which is required for elongation and appears to increase in speed as the egg chamber matures (Cetera et al., 2014). We decreased follicle cell number by knocking down the cell cycle regulator String in the FE over 24 hours (Figure S2D) (Jimenez et al., 1990). Growth and elongation of the egg chamber were unaffected over the stages examined, although gross epithelial defects, including gaps, appear in later stage egg chambers (Figures 2A,B, and S2E). As a positive control for elongation, we disrupted the planar polarity molecule Fat2, which is required for egg chamber rotation (Viktorinová et al., 2009). Fat2-shRNA egg chambers mature without fully elongating, and therefore have a larger average cross-sectional area when compared to wild type egg chambers of the same aspect ratio (Figure 2B). These results demonstrate that egg chamber elongation does not rely on increased follicle cell proliferation, and suggest that the increase in cell number is required to maintain an epithelial covering for the egg chambers as it matures.

**Figure 2:**
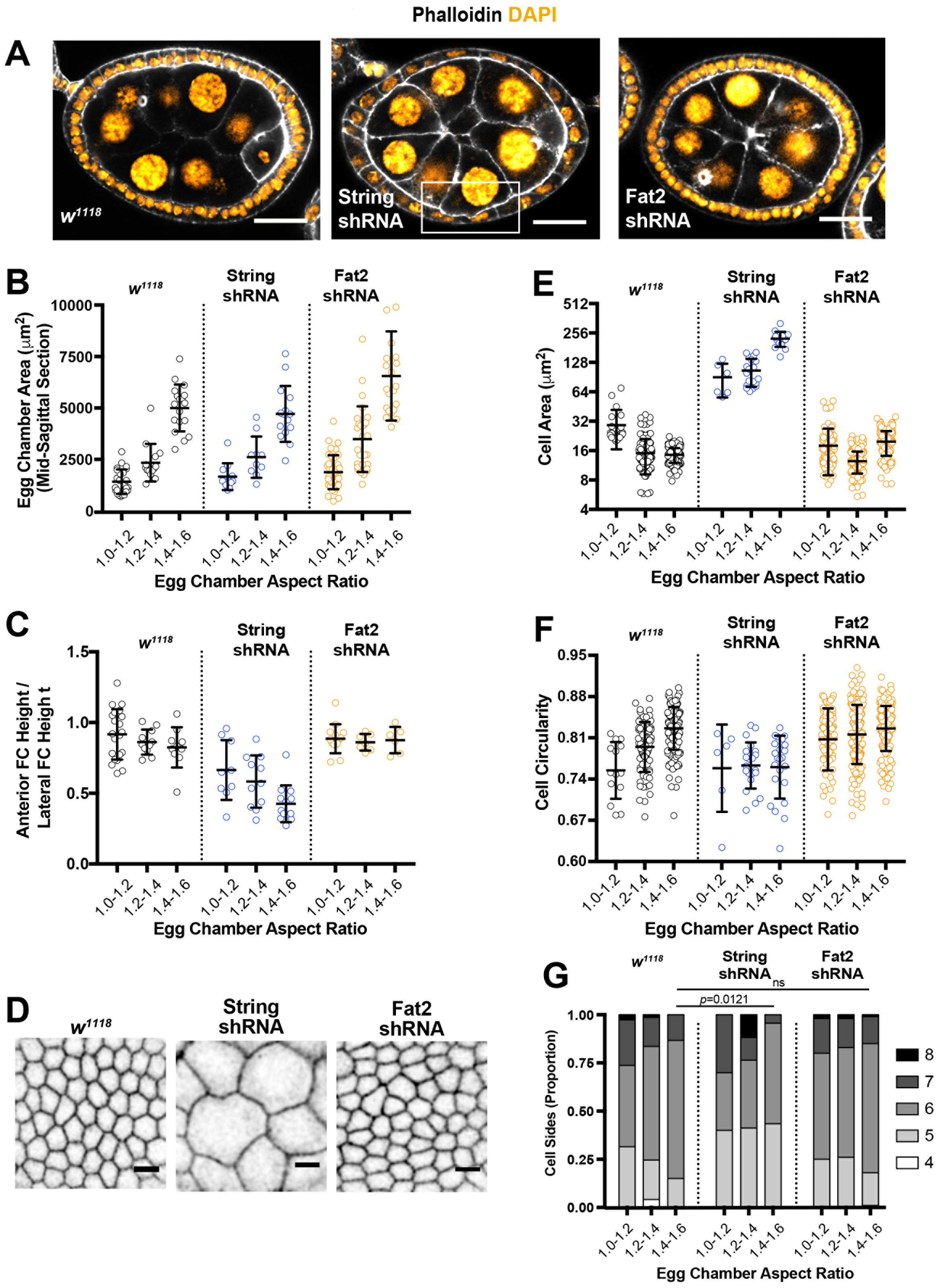
Tissue regularity is a function of cell proliferation. **A and B)** Elongation is unaffected by reduced cell proliferation. String-shRNA expressed for 24 hours alters epithelial shape and cell number (A), but not the relationship between egg chamber aspect ratio and cross-sectional area, which is quantified in (B). Fat2-shRNA is used as a positive control. Egg chambers shown in A have approximately equal widths. Scale bars = 20μm. **C)** Epithelial cell size and shape is affected by decreased proliferation. The epithelial cells in these images are taken from egg chambers with aspect ratios of approximately 1.4. These cells are from different egg chambers than those shown in A. Cell outlines are marked with Basigin::YFP. Scale bars = 5μm. **D, E, F)** Follicle cell area, shape, and polygonality measurements in control, String-shRNA, and Fat2-shRNA egg chambers of different aspect ratios. Cell area in D is shown on a log_2_ scale. These measurements were derived from at least three egg chambers from at least three flies per AR range. Bars represent mean and standard deviation. Statistical significance for cell side distributions was calculated using a chi-squared test. Number of cells used for analysis: *w^1118^* 1.0-1.2 n= 23, 1.2-1.4 n=114, 1.4-1.6 n=160; *mud^3^/mud^4^* 1.0-1.2 n=11, 1.2-1.4 n=191, 1.4-1.6, n=109; String-shRNA 1.0-1.2, n=6, 1.2-1.4 n=19, 1.4-1.6 n=17; Fat2-shRNA 1.0-1.2 n=93, 1.2-1.4 n=167, 1.4-1.6, n=173.

Although the FE in pre-vitellogenic egg chambers remains intact after String depletion, the decrease in cell number alters tissue morphology (Figure 2A). Tissue “height” (length along the apical-basal axis) is a gradient in these egg chambers, with the shortest tissue over the center and the tallest at the anterior and posterior poles (Figures 2A and S2F). The extent of this gradient increases as the egg chamber elongates (Figure 2C). These observations reveal a gradient of tissue tension that is strongest at the middle of the egg chamber, and show that the Stg-shRNA follicular epithelium stretches - like parafilm - to compensate for the decrease in cell number. As in early stage wild type egg chambers, we frequently observed that cell height was uneven in the stretched lateral cells, indicating mechanical stress (Figure 2A, box and Figure S2F). Together, these findings show that elongation causes directional stress on the follicle epithelium, and that this stress is normally relieved by proliferation.

Tension affects follicle cell area as well as height. Follicle cells in Stg-shRNA tissue, where tension is increased, show dramatically increased cross-sectional area relative to wild type (Figures 2D,E). Follicle cell area is also affected by Fat2-shRNA; some Fat2-shRNA egg chambers with aspect ratios between 1.0 and 1.4 are older and larger than their wild type counterparts, and therefore the component follicle cells are smaller (Figure 2E). Fat2-shRNA egg chambers with aspect ratios between 1.4 and 1.6 may have already matured to vitellogenic stages, in which the follicle epithelium stops dividing despite a dramatic increase in egg chamber size. Follicle cells in these egg chambers are larger in area than follicle cells in wild type egg chambers at the same aspect ratio.

These observations show that follicle cells are capable of withstanding substantial mechanical deformation, consistent with observations made in older wild type egg chambers. At stage 9, the post-mitotic epithelium surrounding the anterior nurse cells transitions from cuboidal to squamous, with a dramatic increase in individual follicle cell area. Our results agree with previous work suggesting that this change in morphology can be attributed solely to mechanical stress (Kolahi et al., 2009).

Finally, we measured the effect of reduced proliferation on epithelial cell shape. Measurements of cell circularity and sidedness demonstrate that String-shRNA prevents the follicle epithelium from increasing in regularity as the egg chamber elongates (Figure 2F,G). We conclude that tissue packing regularity in the FE, as in other systems, emerges as a consequence of proliferation (Gibson et al., 2006).

### Interphase Cell Elongation Does Not Predict Division Orientation

Because it is expanding directionally, optimal tissue packing in the FE requires an extra level of proliferation control. One possible mechanism is the elimination of overcrowded cells. However, we and others have failed to observe apoptosis in the FE, with the exception of several (<5) supernumerary polar cells lost by stage 5 (Bergstralh et al., 2015; Besse and Pret, 2003). Another possible explanation is that the rate of proliferation varies across the tissue, but we did not observe a difference in phospho-histone 3 immunoreactivity at different locations in the egg chamber (data not shown).

In a tissue under tension, regularity can be achieved through a bias in the direction of cell division, such that new daughter cells are likely to appear along the stretch axis (Wyatt et al., 2015). Through this mechanism, local tension is relieved (Campinho et al., 2013; Mao et al., 2013; Wyatt et al., 2015). In the *Drosophila* wing disc and pupal notum, and during zebrafish epiboly, localized cortical tension is reflected in cell shape; tension causes interphase cells to deform, giving them a long-cell axis at the apical tissue surface (Bosveld et al., 2016; Campinho et al., 2013; Mao et al., 2013). Interphase deformation has also been observed in a cultured cell system under stretch (Wyatt et al., 2015). In agreement with Hertwig’s Rule, the direction of long-cell axes correlates with planar division orientation, despite the fact that most epithelial cells lose their interphase shape and become round at mitosis (Hertwig, 1884). (By our definition, the orientation of division is the direction in which the daughter cells separate).

We considered the possibility that interphase cell shape determines cell division orientation in egg chambers. We found that follicle cell shapes are not oriented in the direction of tissue elongation but rather show a significant bias against it (Figures 3A,B). Furthermore, the extent of individual cell elongation correlates strongly with this directional bias (Spearman rank correlation test, p = 0.0003) (Figure S3A). Cell shape is attributed to the collective cell migration that drives egg chamber rotation, which requires a traction force on basal cell surfaces (Cetera et al., 2014; Duhart et al., 2017b; Haigo and Bilder, 2011; Viktorinová et al., 2017). This force appears to deform the entire cell, since the cell shape bias is the same at the apical and basal cell surfaces (Figure 3B).

**Figure 3:**
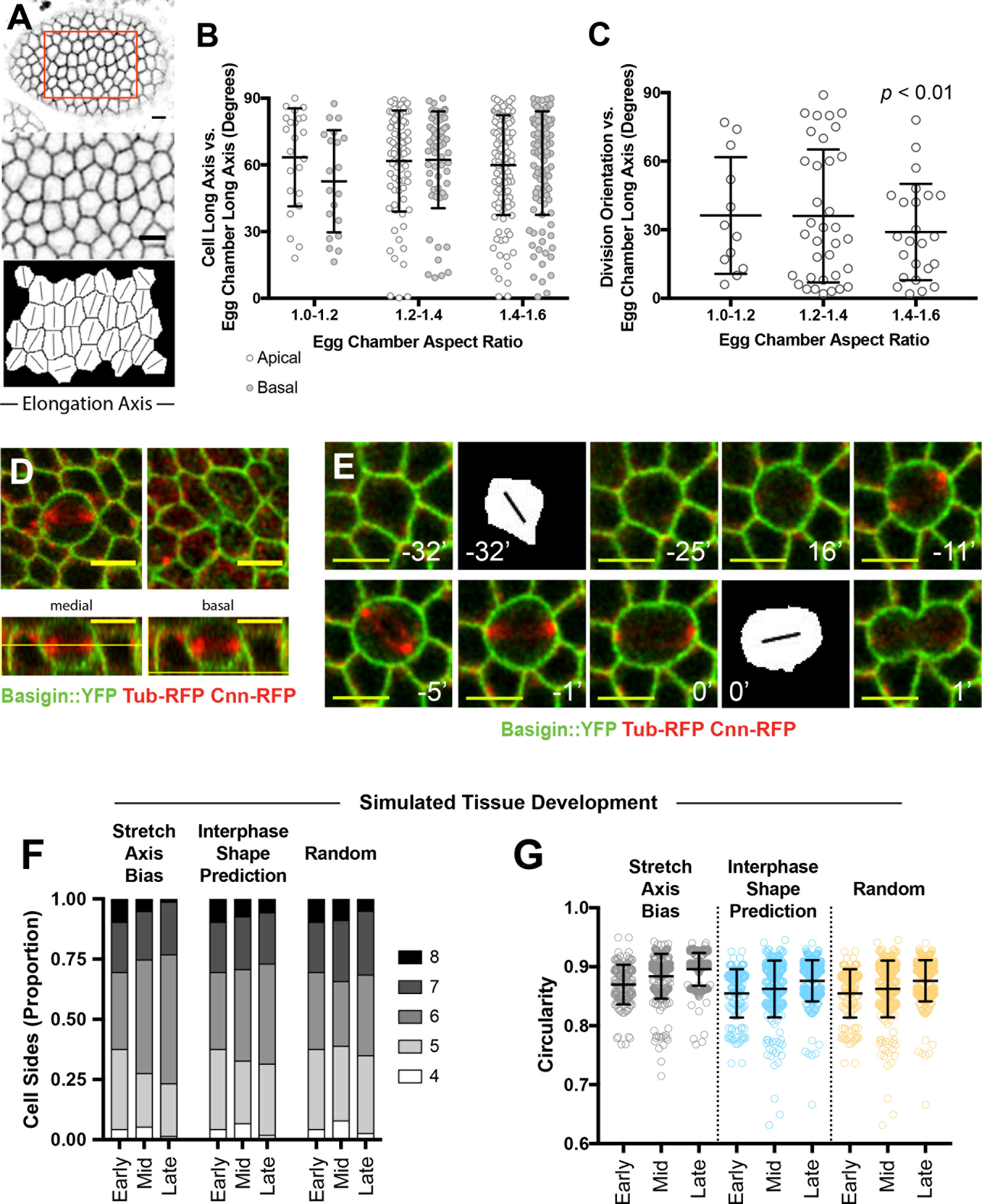
Interphase shape does not predict division orientation in the follicular epithelium. **A and B)** Follicle cell elongation is biased perpendicular to the elongating axis of the egg chamber. Long-axis and division angles are presented as rose plots for ease of comparison. In A, segmentation of an egg chamber with an aspect ratio of 1.5 reveals that follicle cell elongation is usually perpendicular to the elongation axis of the egg chamber. Cell outlines were marked with Bsg::YFP. Error bars = 5μm. Quantification of cell elongation angles is presented in B. (1.0-1.2 n=23, 1.2-1.4 n=76, 1.4-1.6 n=114.) **C)** Live cell imaging shows that planar division orientation is initially random in the follicular epithelium, with a significant bias towards the egg chamber elongation axis evident in egg chambers with aspect ratios between 1.4 and 1.6. (Number of cells: 1.0-1.2 n=12, 1.2-1.4 n=35, 1.4-1.6 n=24.) **D)** Neighboring cells can form new attachments underneath (basal to) a mitotic cell that has moved apically. Scale bars = 5μm. Thin yellow lines in the XZ reconstructions indicate the z-position of the two XY focal planes. **E)** Live imaging shows that the long axis of a follicle cell at interphase does not predict the orientation of division. Long axis orientations at the two timepoints shown were determined as in A. **F and G)** Sidedness and circularity are affected by planar division orientation in a simulated epithelium. One round of doubling extending over a six-hour period was simulated in a stretched tissue under different conditions for division orientation. The three time points shown reflect cell geometry in the first two hours (early), second two hours (mid), and last two hours (late).

We would expect apical shape to have a stronger impact on cell division for two reasons. First, mitotic cells in the follicular epithelium frequently pull away from the basement membrane before dividing, and we show here that these detached cells can be physically excluded from the basal surface by neighboring follicle cells, which adjust to form attachments underneath them (Figure 3D) (Bergstralh et al., 2015). Second, planar orientation in the wing disc, pupal notum, and embryonic mesectoderm is determined by cortical tension at the apical tissue surface (Bosveld et al., 2016; Mao et al., 2013; Wang et al., 2017).

Perpendicular bias in the orientation of cell long-axes relative to the A-P axis presents a difficulty for the tissue, since corresponding cell divisions would be expected to expand the tissue contrary to the direction of egg chamber elongation. This problem is resolved by the finding that planar cell division orientations do not correspond to the overall bias in cell long-axes. Divisions are randomly oriented within the planar axis until later elongation (AR 1.4-1.6), at which point the average angle demonstrates a significant bias from random (Wilcoxon Signed Rank Test, *p* = 0.0011) towards the direction of tissue elongation (Figure 3C). We consider the average planar angle that we measured (approximately 30°) to reflect the directional bias in tissue expansion, since the tissue at this point is approximately two-thirds as wide as it is long. We measured division axes by the position of centrosomes at anaphase or early telophase, and performed the measurements shown here in live tissue. Fewer divisions were imaged in the longer egg chambers, which is likely because these are mostly at stage 6, when division ceases.

Live imaging also revealed divisions that violate the long axis prediction (Figure 3F and Supplementary Movie 1). In nine complete divisions measured starting from interphase, the average difference between the interphase long axis and the division angle was 35.2°, with a standard deviation of 24.8°.

Planar cell division orientation is likely to be most important in longer egg chambers, which are under the greatest tension. To explore this possibility, we built a computational model for cell division in a stretched tissue. Because follicle cell division ends in longer egg chambers, we simulated only one cell doubling. We tested three conditions: division biased along the stretch axis, 30° off the stretch axis (reflecting the difference between observed spindle angle and the interphase shape prediction), and random. Cell areas decreased by roughly half, in agreement with a rate of proliferation that outpaces tissue expansion (Figure S3B). We observed that division promoted optimal packing (hexagonality) in the stretch bias condition, and this effect was disrupted by division misorientation (Figure 3F). Cell circularity was impacted to a lesser extent (Figure 3G). We conclude that follicle cells break Hertwig’s Rule to promote tissue regularity.

### Distinct Mechanisms Control Spindle Orientation in the Apical-Basal and Planar Axes

These findings raise the question of how planar orientation is controlled. Given that divisions in longer egg chambers are biased towards the elongating (stretch) axis of the tissue, we asked whether cortical tension, which is transmitted through apical junctions, influences division orientation. During mitosis, cortical tension should have a stronger influence on the cell than basal traction, since mitotic cells lose contact with the basement membrane (Figure 2C).

We showed previously that the canonical Pins/Mud machinery orients follicle cell spindles in the A-B axis, but did not address the possibility that it also participates in planar division orientation (Bergstralh et al., 2013). In the imaginal wing disc and pupal notum, Mud localizes to vertices (junctions between three or more cells) throughout the cell cycle (Bergstralh et al., 2016; Bosveld et al., 2016). This pattern results in an anisotropic distribution of the pulling force during mitosis, and translates the interphase long cell axis (at the apical surface) into planar division orientation (Bosveld et al., 2016). The position of cell vertices at interphase can therefore be used to predict division orientation in the wing disc/pupal notum (Bosveld et al., 2016). We found that the distribution of cell vertices does not correspond to division orientation in the FE (Figure S4A). This observation is explained by the composition of cell vertices in these tissues. The wing disc and pupal notum are considered “mature,” since they maintain specific protein complexes called tricellular junctions (TCJs) at the level of septate junctions at cell vertices. TCJs are necessary for Mud localization (Bosveld et al., 2016). We found that the FE is immature, since it does not demonstrate immunoreactivity to the TCJ marker Gliotactin (Figure S4C). We examined Mud and Pins localization in the FE and another immature epithelium, the early embryonic neuroectoderm (stages 8-9), and observed that both proteins localize around the entire mitotic cell cortex (Figure 4A,B and S4B) (Schulte et al., 2003). Live imaging in these tissues suggests that spindle orientation in the A-B and planar axes are distinct, since we observed extensive rotation of the spindle in the plane even after A-B orientation had been achieved (once both poles appear in the tissue plane) (Figure 4C and Supplementary Movies 2,3). Similar spindle rotation within the tissue plane is observed during zebrafish epiboly (Larson and Bement, 2017). Contrastingly, less planar rotation is observed in the imaginal wing disc, as would be expected from an anisotropic pulling force that orients the spindle in the A-B and planar axes concomitantly (Figure 4D).

**Figure 4:**
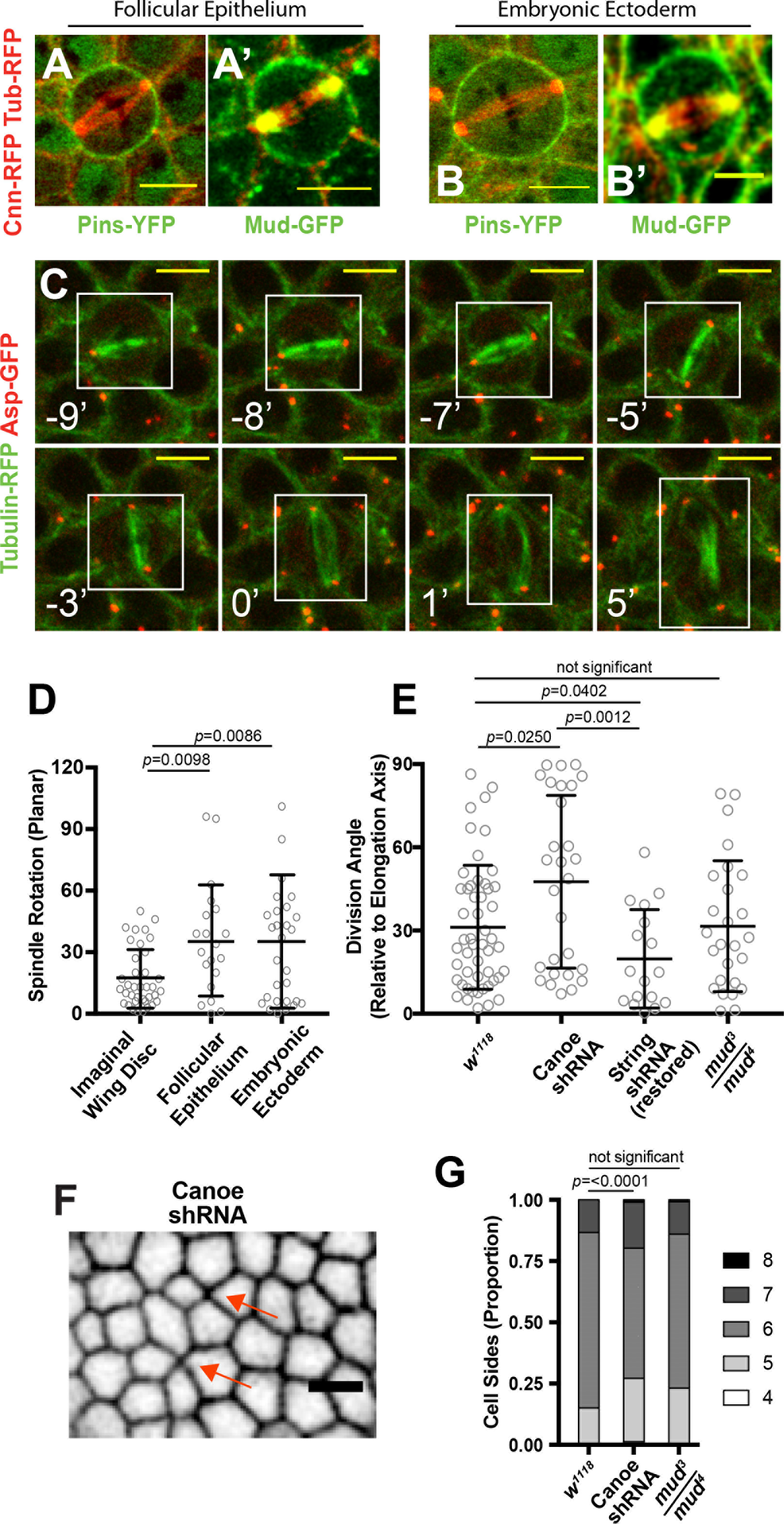
Cortical tension drives division orientation independently of Pins/Mud. **A and B)** Pins and Mud are symmetric around the cell cortex in mitotic follicle cells (A and B) and mitotic cells in the embryonic neurectoderm (A’ and B’). C) Live imaging reveals a mitotic spindle rotating in the plane of the epithelium until anaphase at timepoint 0. **D)** Mitotic spindles in the follicle epithelium and embryonic neurectoderm undergo more rotation in the plane of the tissue than spindles in the imaginal wing disc (unpaired t-test with Welch’s correction). Number of cells used for analysis: Imaginal wing disc n=37; Follicular epithelium n=20; Neuroectoderm n=30. **E)** Planar division orientation in the follicular epithelium relies on cortical tension but is independent of the spindle orienting machinery. Division angles were calculated relative to the elongating axis of the egg chamber. Statistical significance was determined by Mann Whitney U. This analysis includes data combined from live and fixed egg chambers. *w^1118^* n=27 fixed, 24 live; Canoe-shRNA n=28 fixed; String-shRNA (restored) n=17 fixed; *mud^3^/mud^4^* n=20 fixed, 5 live. **E, F)** Tissue topology is affected by disruption of Canoe. E is a representative image of a Canoe-shRNA follicle epithelium. Cell outlines are marked with Basigin::YFP. This egg chamber has no mitotic cells and and AR of 1.58. The arrows point to four-cell junctions, which are unusual at this stage. Scale bar = 5μm. Sidedness for egg chambers with aspect ratios 1.4-1.6 is quantified in F. Statistical significance for cell side distributions was calculated using a chi-squared test. *w^1118^* 1.0-1.2 n=38, 1.2-1.4 n=73, 1.4-1.6 n=284; Canoe-shRNA 1.0-1.2 n=35, 1.2-1.4 n=98, 1.4-1.6 n=310; *mud^3^/mud^4^* 1.0-1.2 n=11, 1.2-1.4 n=135, 1.4-1.6 n=251.

In a cultured cell model, cortical tension is translated into mitotic spindle orientation by the adherens junction component E-cadherin, which can interact directly with LGN (vertebrate Pins) (Hart et al., 2017). Stress causes E-cadherin to become locally enriched during interphase, and this enrichment results in polarized recruitment of LGN to the cortex at mitosis. We considered the possibility that either Shotgun (*Drosophila* E-Cadherin) or Canoe (*Drosophila* Afadin), which is also at adherens junctions and is reported to directly regulate the Pins/Mud machinery in cultured cells and asymmetrically dividing *Drosophila* cells, could perform a similar role in the FE (Carminati et al., 2016; Johnston et al., 2013; Speicher et al., 2008; Wee et al., 2011). However, our data do not support such a mechanism. First, adherens junctions are not proximal to spindle poles in the FE (Figure S4D) (Bergstralh et al., 2013). Second, Pins and Mud are symmetric with respect to the tissue plane at mitosis (above). Third, we did not observe planar polarized enrichment of either Shotgun or Canoe in interphase follicle cells (Figures S4E,F). Finally, planar division angle is unaffected in *mud*^*3*^/*mud*^*4*^ egg chambers (Figure 4E). These findings, combined with those outlined above, do not support a model whereby planar division orientation is determined by adherens junction-mediated regulation of the Pins/Mud machinery. We also found that spindle orientation in the A-B axis is normal in mitotic clones mutant for the null allele *cno*^*R2*^, ruling out a role for Canoe in regulating that aspect of division orientation (Figure S4G). Together, these results show that division orientation in the planar axis is independent from the Pins/Mud machinery, but do not rule out a role for cortical tension.

We tested the possibility that cortical tension governs planar orientation by using temperature modulation of String-shRNA activity to increase tension on dividing cells. The UAS-Gal4 transcription system is efficient at 29°C, but not 18°C. We therefore depleted String at 29°C for 24 hours, as in Figure 2, to increase tension, then restored String activity by incubating the flies at 18°C for 24 hours. Spindle orientation in fixed tissue required three-dimensional reconstruction, as described in the Methods. Although we observed aberrant spindles in the flattest cells (not shown), consistent with prior work, we measured planar division angles in the remaining cells in egg chambers with AR 1.4-1.6 (Lancaster et al., 2013). These cells normally show an average planar orientation of 31° off the elongation axis. The planar division angle bias towards the elongating axis in the String-shRNA (restored) condition was stronger, with an average of 20° (Figure 4E).

Adherens junctions are responsible for mechanotransduction and mechanosensing in epithelial cells (reviewed in (Lecuit et al., 2011)). We therefore measured planar division angle after disrupting Canoe. Although it does not have a role in A-B spindle orientation (above), we hypothesized that Canoe/Afadin could impact planar orientation because it is part of the machinery that links the actomyosin cytoskeleton to adherens junctions in epithelial cells (Sawyer et al., 2011; Sawyer et al., 2009). While Afadin/Canoe is not required to maintain tissue tension, it should be expected to transmit tissue tension to the cell (Choi et al., 2016). Consistent with this, Canoe-shRNA follicle cells in egg chambers with AR 1.4-1.6 have an average planar division angle that is not statistically different from random (Wilcoxon Signed Rank Test) and is higher than wild type (Figure 4E). (Immunostaining was used to confirm that Canoe-shRNA depletes Canoe protein expression (Figure S4H).) This effect appears to be at the cellular, rather than tissue level, since Canoe-shRNA did not affect tissue elongation (Figure S4I).

We next examined the impact of Canoe-shRNA on tissue topology. Consistent with the computational model (Figure 3), cell size and circularity were not appreciably different after Canoe disruption (Figures S4J,K). However, follicle cells in egg chambers with aspect ratios 1.4-1.6 were less likely to be hexagonal than in wild type (Figures 4E,F). We conclude that tension-driven division orientation contributes to regular tissue packing.

We have shown that tissue regularity in a proliferating epithelium requires cells to divorce their division orientation from their interphase cell shape. Although previous work shows that planar division orientation relies on anisotropy of the Pins/Mud machinery, we show that orientation along the plane of the tissue is independent of Pins/Mud. Instead, it relies on Canoe, which links actomyosin-based cortical tension to the cell. These findings show that division orientation is bimodal, and occurs independently in the apical-basal and planar tissue axes (Figure 5). However, the molecular mechanism that translates tension into division orientation remains to be elucidated. Our study also raises the question of whether bimodal spindle orientation is special to the FE, or a broad feature of immature epithelia. Future work will address this problem.

**Figure 5:**
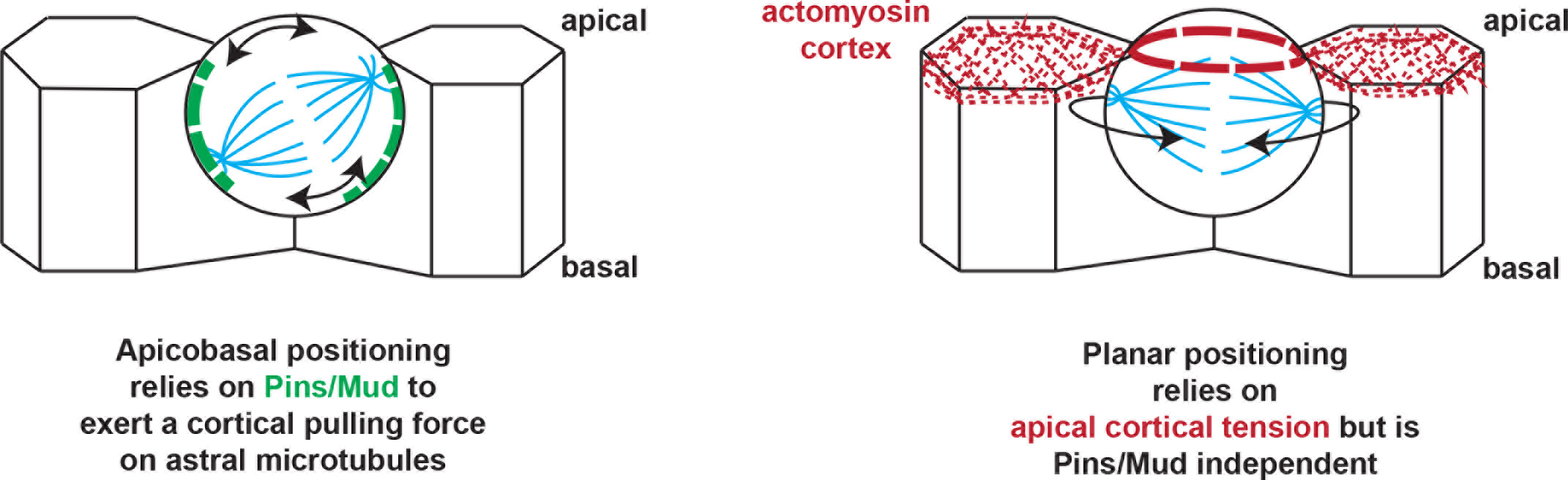
A model for division orientation in the follicular epithelium. Orientation is achieved separately in the apical basal and planar axes. Pins/Mud in green, Apical
actomyosin in red, microtubules in blue.

## Materials and Methods

***Drosophila* mutants and marker stocks**: The following mutant alleles and transgenic constructs have been described previously: *mud*^*3*^ and *mud*^*4*^ (Yu et al., 2006), *cno^R2^* (Sawyer et al., 2009), Ubi-α-Tub-RFP (Basto et al., 2008), Ubi-α-Tub84B-GFP (Rebollo et al., 2004), Ubi-Abnormal spindles-GFP (Rujano et al., 2013), Ubi-Centrosomin-RFP (Basto et al., 2008), Ubi-Centrosomin-GFP (Conduit et al., 2010), Ubi-Pins-YFP, *pins*^*p62*^ (David et al., 2005), Basigin::YFP (Bergstralh et al., 2015; Lowe et al., 2014), Mud-GFP (Bosveld et al., 2016), Pavarotti-KLP::GFP (Minestrini et al., 2003), Shotgun::GFP (Huang et al., 2009) and Spaghetti squash::mCherry (Martin et al., 2009). We thank the Transgenic RNAi Project at Harvard Medical School (NIH/NIGMS R01-GM084947) for UAS-Cno-shRNA (HMS00239), UAS-String-shRNA (HMS00146) and UAS-Fat2-shRNA (HMS02136). The following background stocks were used to generate mitotic clones, which were induced by heat shock at 37° for multiple periods of two hours: RFP-nls, hsflp, FRT19A and hsflp; FRT40A RFP-nls, and hsflp;; FRT82B RFP-nls. Ectopic expression in follicle cells was driven by Traffic Jam-Gal4 (Olivieri et al., 2010), or actin5c-FLPout-Gal4, UAS-GFP (Fig. S4G only). For the String-shRNA experiments, flies developed to adulthood at 18°C, and were transferred to 29°C for 24 hours. For the Canoe-shRNA and Fat-shRNA experiments, flies were incubated at 29°C for 48 hours.

### Reagents

The following antibodies were used in this study: rabbit anti-Canoe (gift from M. Peifer) (Sawyer et al., 2009), rabbit anti-Centrosomin (gift from J. Raff) (Lucas and Raff, 2007), rabbit anti-Mud (gift from R. Basto) (Rujano et al., 2013), rabbit anti-Pins (gift from F. Matsuzaki) (Izumi et al., 2006), rabbit anti-Gliotactin (gift from V. Auld), mouse anti-Armadillo (Developmental Studies Hybridoma Bank, N2 7A1 ARMADILLO, 12/10/15), and mouse FITC-conjugated anti-a-tubulin (Sigma, clone DM1A, Lot#114M4817V). Conjugated secondary antibodies were purchased from Thermo Fisher Scientific. Phalloidin was purchased from Invitrogen and Vectashield with DAPI was purchased from Vector Labs. Primary and secondary antibodies were used at a concentration of 1:150.

### Imaging

Live cell imaging was performed in 10S halocarbon oil (egg chambers), or Schneider’s medium (wing discs), as previously described (Bergstralh et al., 2015). Both live-and fixed-tissue imaging were undertaken on a Leica SP5 (63x/1.4 HCX PL Apo CS Oil). For live imaging in three dimensions, Z-stacks of planes spaced 0.5μm apart were taken at one-minute intervals. Image collection and processing (Gaussian blur) were performed with Leica LAS AF and ImageJ, respectively.

### Topology Measurements

The long and short axes of the egg chamber were determined using the EllipseFitter macro for ImageJ. Apical and basal surfaces of the follicle cells were manually identified from acquired Z-stacks. Follicle cell membranes were segmented using a custom Java macro written for use in ImageJ which included use of features from the Biovoxxel Toolbox for ImageJ (Brocher, 2015). Cells at the edge of the field of view were excluded from analysis. The area, circularity, aspect ratio and the angle of the cell long axis were calculated using the Analyze Particles tool. The cell long axis is identified in this method by fitting an ellipse of best fit, with the Angle result giving the angle between the primary axis and a line parallel to the x-axis of the image, which was manually set to correspond to the AP axis of the egg chamber. The angle was then corrected to be between 0-90°. Circularity was determined by:

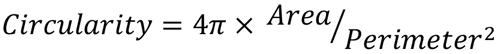

The number of cell sides (vertices) was identified using the ‘Neighbor analysis’ macro from the Biovoxxel Toolbox using watershedding and Voronoi analysis. All of these measurements are based on at least three egg chambers, from three flies, per aspect ratio. Centrosome angles with respect to the long axis were calculated using Image J. Statistical analyses were performed using Prism (GraphPad).

### Circumferential Cell Number and Tissue Height Analysis

These measurements were performed manually using ImageJ. The analyst was blinded to tissue genotype. Lateral FC height is an average of measurements at two opposing positions midway through the egg chamber.

### Division Angle Analysis (Live Tissue)

The long axis of the egg chamber was determined using EllipseFitter for ImageJ. Division angles were measured by drawing a line between the two centrosomes and determining the angle between this line and the long axis.

### Vertex Distribution Analysis (Live Tissue)

Images were segmented and the position of cell barycenters, vertices and the vectors describing the position of the cell vertices from the cell barycenters were identified using a custom MatLab script. The vector describing the long axis of the cell as described by the vertex positions (‘Vertex distribution’) was then calculated as previously described (Bosveld et al 2016). We also considered the possibility that the distance of vertices from the barycenter could influence the magnitude of the pulling force, and therefore calculated the angle of the average of the summed vectors that describes the position of the vertices (‘Vertex vector’).

### Division Angle Analysis (Fixed Tissue)

Division angles were determined using algorithms custom written in Python3. Qualitatively, the spindle was rotated about the long axis of the egg chamber to produce a top down view centered on the midpoint of the spindle. The angle of division was then measured as the deviation of the spindle orientation from the long axis. Mathematically, images were first thresholded and segmented in each plane of a z-stack based upon the tubulin staining to create binary image stacks of the egg chamber. The long axis of the egg chamber was calculated using a covariance matrix of the second central moments of the thresholded images to determine the unit eigenvectors, which represent the directions of the volume’s principal axes. The axis corresponding to the largest eigenvalue was taken as the long-axis of the egg chamber. To determine spindle orientations, centrosome positions in x, y, and z were manually recorded. The midpoint between each centrosome pair was calculated, and a line was drawn from the midpoint to the point where the long axis intersected with the edge of the egg chamber. From this, a unit vector was calculated which pointed from the spindle midpoint to one end of the chamber. A plane of projection was defined by the orthogonal directions of the cross-product of this unit vector with that of the long-axis, and the long-axis itself. Each centrosome pair was projected onto this plane using a dot product between the spindle unit vector and the unit vectors defining the plane of projection, yielding the desired top-down view. The angle of deflection between the projected centrosome unit vector and the long-axis was calculated by taking the inverse tangent of the two components of the projected spindle unit vector. These analyses were performed by researchers (PWO and AVS) blind to egg chamber genotype. For consistency with live imaging, divisions at the egg chamber termini were excluded.

### Computational Model

We simulated a vertex model of the growing follicular epithelium (Fletcher et al., 2014). For simplicity, we neglected curvature and instead simulated a two-dimensional planar tissue, imposing doubly periodic boundary conditions since the tissue is an edgeless epithelium. The model was non-dimensionalised such that all lengths and times were scaled with a typical cell diameter by one hour, respectively. Assuming overdamped motion in this model, the position ***r***_*i*_(*t*) of each vertex *i* (where three cells meet) evolves according to the equation of motion

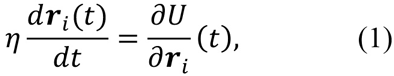

where η = 0.04 denotes a drag coefficient and the ‘free energy’ function U is given by (Farhadifar et al., 2007)

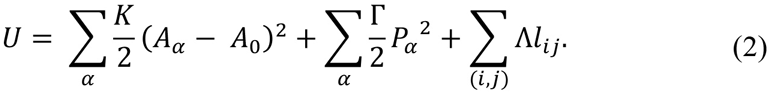

This term includes contributions from cellular bulk elasticity, cortical contractility, and the combined effect of cell-cell adhesion and junctional contractility. Here, A_α_ and P_α_ denote the apical surface area and perimeter of cell α, A_o_ = 1 is a common ‘target’ area, l_ij_ is the length of the edge shared by vertices i and j, and the coefficients K = 1, Γ = 0.04, and Λ = 0.12 together govern the strength of the individual energy contributions.

We simulated the growth of the egg chamber by imposing a slow increase in the width and length of the periodic domain, with speeds *s*_*x*_ = 0.001 and 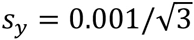. For simplicity, we assumed that cell cycle times are independent and randomly drawn from the uniform distribution with minimum *τ*_*min*_ = 3 and maximum *τ*_*max*_ = 5. When a cell is ready to divide, we created a new cell-cell interface at a specified angle *θ* to the *x*-axis through the centroid of the parent cell. We considered three alternative choices for the angle of division: (1) biased towards the stretch axis of the tissue, such that *θ* is drawn from the wrapped normal distribution with mean 30 degrees and variance 10 degrees; (2) randomised division, such that *θ* is drawn from the uniform distribution with minimum 0 degrees and maximum 180 degrees; (3) orientation perpendicular to the long axis of the parent cell (stretch axis bias).

Each simulation started from an initial cell packing generated by computing the Voronoi tessellation of 36 seed points distributed uniformly at random in a square of width *L*_0_ = 6, and relaxing this configuration to mechanical equilibrium. The tissue was then evolved over discrete time steps of length Δ*t* = 0.0001. At each time step we: updated the age of each cell and implemented any divisions; performed a neighbour exchange (T1 transition) for any cell-cell interface whose length dropped below *d*_*min*_ = 0.01, replacing it with a new orthogonal interface of length 1.5*d*_*min*_; removed any cell (T2 transition) whose area dropped below *A*_*min*_ = 0.001; used an explicit Euler method to integrate equation (1) and updated the positions of all vertices simultaneously; and updated the width and length of the periodic domain. Simulations were implemented in the software Chaste (Fletcher et al., 2013; Mirams et al., 2013).

## Acknowledgements

We are grateful to Holly Lovegrove and David Bilder for their useful comments during our study. We thank Tyler Wilson and Allison Bhattacharya for technical assistance.

## Author Contributions

DTB conceived the project and designed the experiments. AGF designed the computational model and performed and analyzed simulations. AVS and PWO designed the algorithms for fixed-tissue division analyses. DN, NSD, and DTB performed the imaging. TMF and DTB designed the analysis and together with AVS and PWO analyzed the images. TMF and DTB wrote the manuscript.

---

**Supplementary Figure 1.**
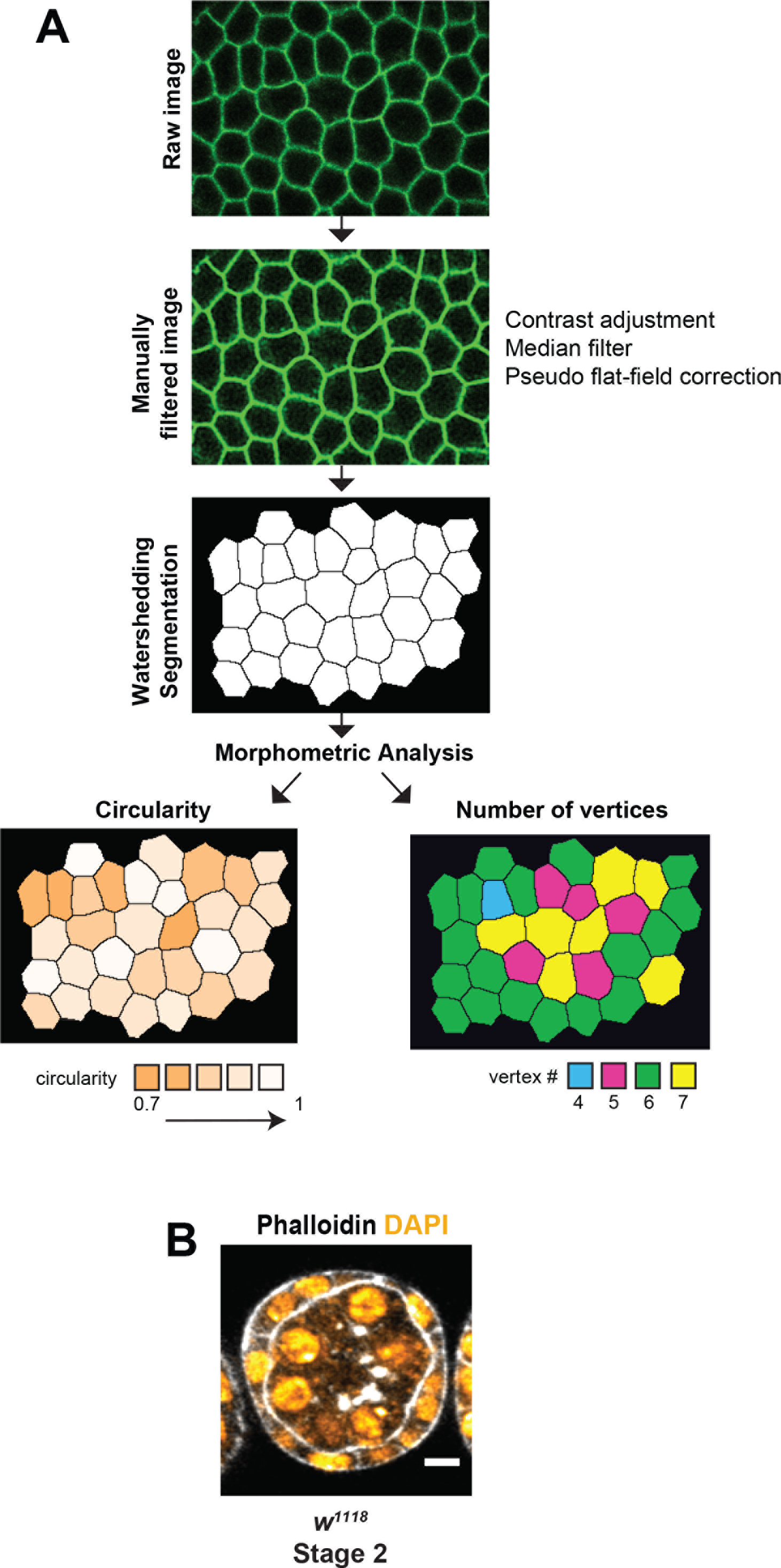
**A)** Visual workflow of image analysis. **B)** Irregular cell height in an early stage egg chamber. Scale bar = 5μm.

**Supplementary Figure 2.**
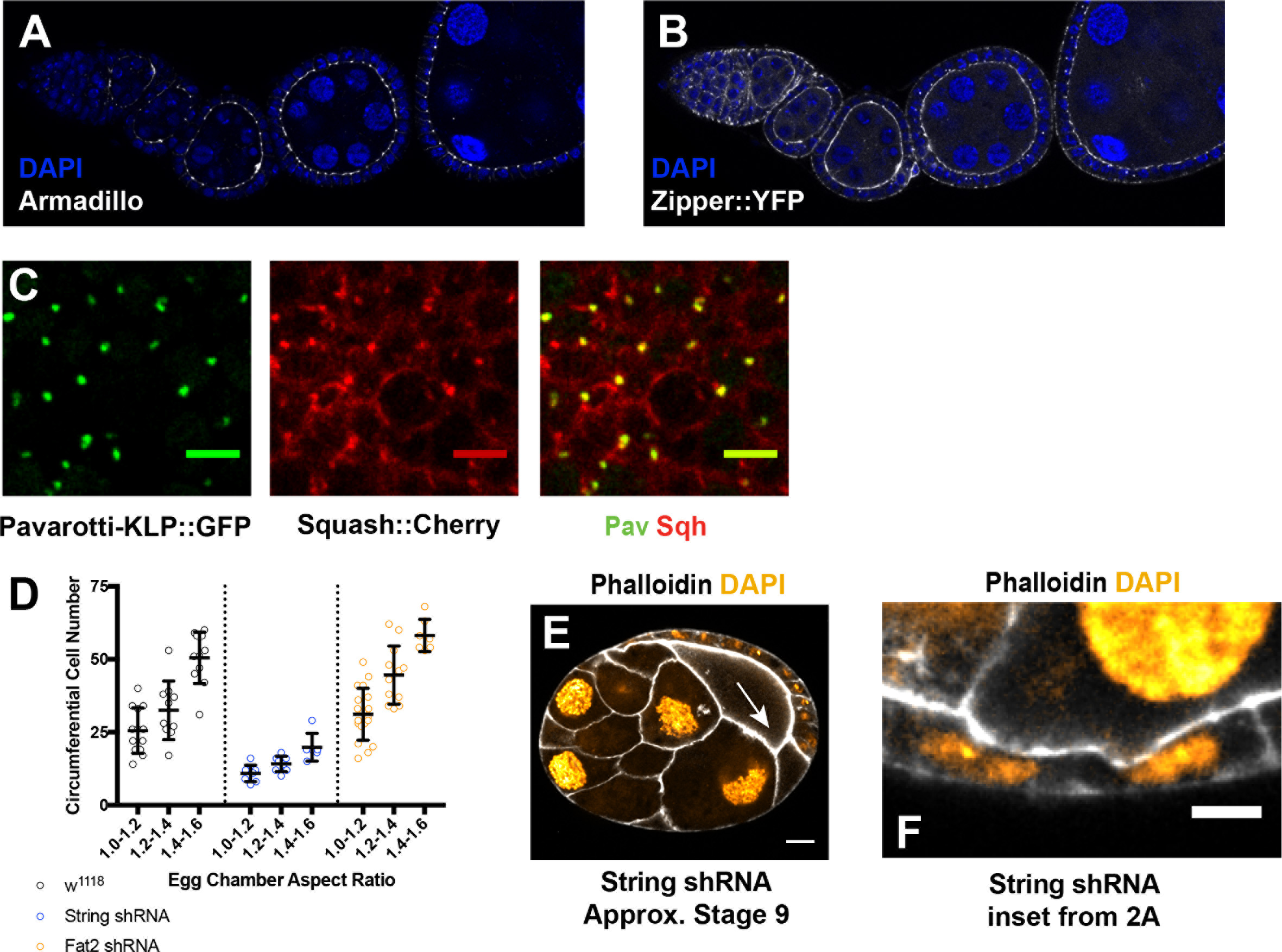
**A and B)** Expression of Armadillo (A) and Zipper (B) in follicle cells does not increase as the egg chamber elongates. **C)** “Blobs” of Squash::Cherry at the medial cell cortex usually colocalize with the ring canal marker Pavarotti-KLP::GFP. **D)** The number of cells around the circumference in egg chambers of increasing aspect ratio is altered by disruption of either String (blue) or Fat2 (orange). Bars represent mean and standard deviation. *w^1118^* 1.0-1.2 n=13, 1.2-1.4 n=10, 1.4-1.6 n=11; String-shRNA 1.0-1.2 n=8, 1.2-1.4 n=8, 1.2-1.4 n=6; Fat2-shRNA 1.0-1.2 n=19, 1.2-1.4 n=12, 1.4-1.6 n=7. **E)** Tissue defects, including gaps (arrow), are observed in later stage egg chambers after String depletion. **F)** The ratio of follicle cell height at the lateral midpoint of the egg chamber to cell height at anterior of the egg chamber is altered by disruption of either String (blue) or Fat2 (orange). Bars represent mean and standard deviation. *w^1118^* 1.0-1.2 n=21, 1.2-1.4 n=11, 1.4-1.6 n=10; String-shRNA 1.0-1.2 n=9, 1.2-1.4 n=11, 1.2-1.4 n=13; Fat2-shRNA 1.0-1.2 n=14, 1.2-1.4 n=7, 1.4-1.6 n=6. **G)** Inset from Figure 2A shows irregular height in a laterally-positioned follicle cell after String depletion. Scale bar = 5μm.

**Supplementary Figure 3.**
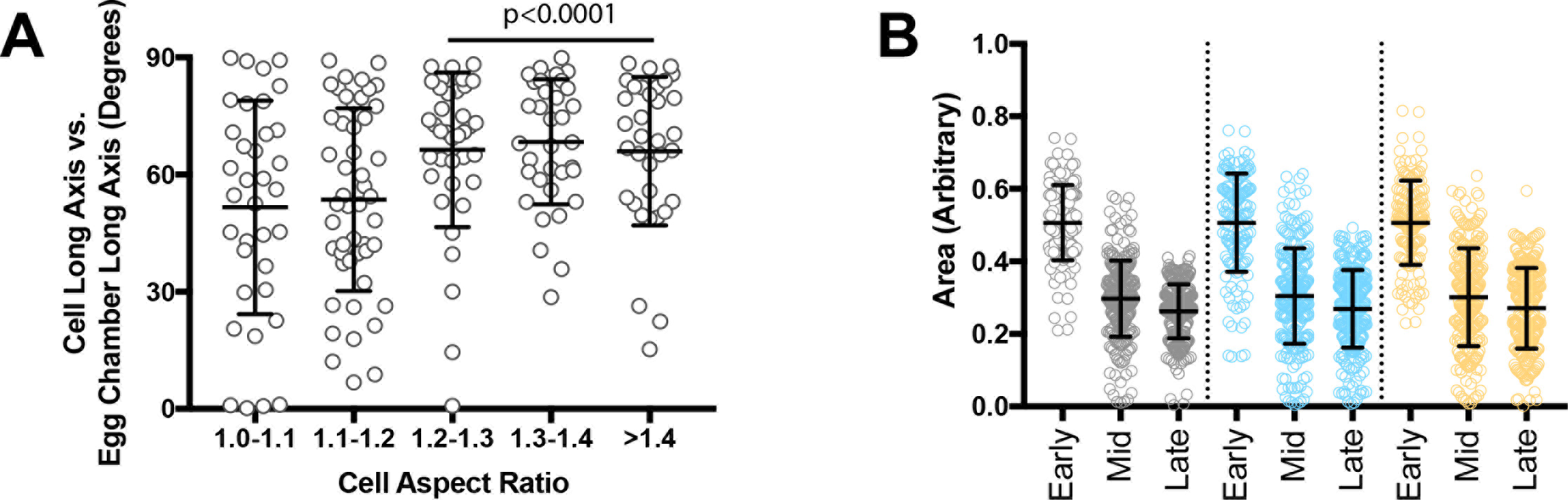
**A)** Interphase cell elongation is biased away from the elongating axis of the egg chamber. Number of cells: 1.0-1.2 n=23, 1.2-1.4 n=76, 1.4-1.6 n=114. **B)** The orientation of follicle cell long axes correlates with the extent of follicle cell elongation (cell aspect ratio). Put another way, the longer the cell, the more likely it is to be oriented perpendicular to the elongating axis of the egg chamber. In this graph, cells quantified in C were binned into elongation groups to facilitate comparison. Average angles in the three longest groups differ from random (Wilcoxon Signed Rank Test). Correlation between long axis and spindle angle is also observed, as described in the text. **C)** Number of cells: 1.0-1.2 n=12, 1.2-1.4 n=35, 1.4-1.6 n=24. Apical elongation axes and cell division axes are also represented in rose plots in Figure 3B and C, respectively, and are shown here to illustrate the distribution and number of data points. Bars represent mean and standard deviation. **D)** Average cell size decreases by half after one round of doubling over six hours in a simulated epithelial tissue undergoing stretch. Cell size in this simulation is not affected by the planar orientation of division. The three time points shown reflect sizes in the first two hours (early), second two hours (mid), and last two hours (late).

**Supplementary Figure 4.**
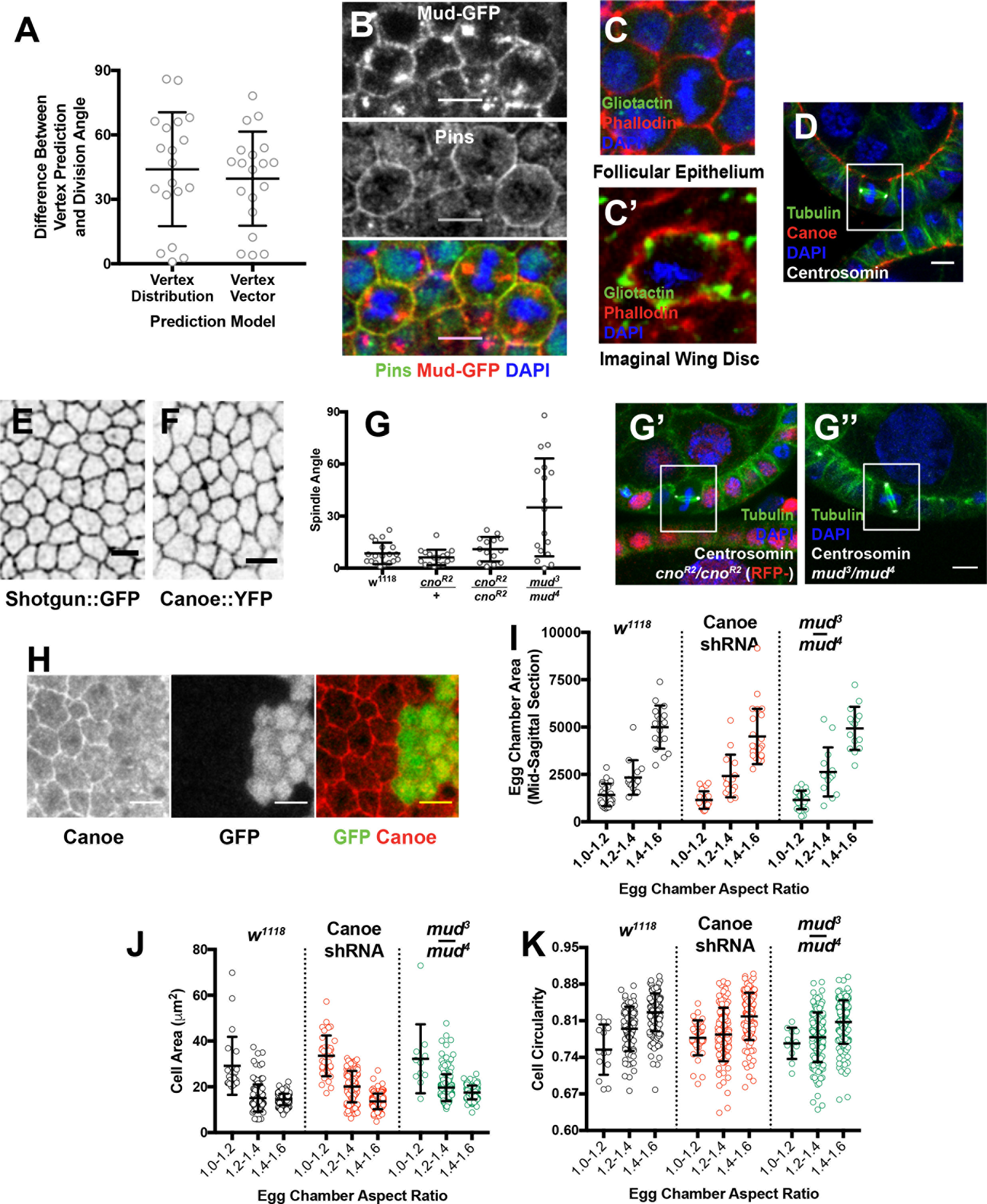
**A)** The vertex long axis of cells does not correlate with the division angle. Neither the cell vertex angular distribution nor the vector sum that describes the distribution of vertices from cell barycentres correlates with the division angle in metaphase cells. Number of cells n=19. **B)** Fixed images show that Pins and Mud-GFP colocalize around the cell cortex in mitotic follicle cells. These results confirm the live imaging shown in Figure 4B. Scale bar = 5μm. **C)** Gliotactin immunoreactivity is not observed in the FE. The imaginal wing disc (C’), in which Gliotactin localizes to cell vertices, is used as a control. **D)** The adherens junction protein Canoe is not proximal to the spindle in mitotic follicle cells. The mitotic cell is outlined in the white box. Scale bar = 5μm. **E and F)** Neither Shotgun (E) nor Canoe (F) show obvious planar polarized enrichment in follicle cells. Scale bars = 5μm. **G)** Apical-basal spindle orientation is not disrupted in mitotic clones mutant for the null allele *cno^R2^*. *mud^3^/mud^4^* egg chambers are used as a positive control. J’ and J’’ show representative spindles in *cno^R2^* and *mud^3^/mud^4^* mutant cells. The mutant clone in J’ is marked by the absence of nls-RFP. Scale bars = 5μm. **H)** Canoe-shRNA depletes Canoe expression in the FE. Canoe-shRNA was driven in a mitotic clone, which is marked by GFP. Canoe immunoreactivity is not observed in the clone. **I and J)** Follicle cell size and circularity measurements in control, Canoe-shRNA, and *mud^3^/mud^4^* egg chambers of different aspect ratios. These measurements were derived from at least three egg chambers from at least three flies per AR range. Bars represent mean and standard deviation. w^1118^ 1.0-1.2 n=11, 1.2-1.4 n=139, 1.4-1.6 n=161; String shRNA 1.0-1.2 n=6, 1.2-1.4 n=21, 1.2-1.4 n=23; Fat2 shRNA 1.0-1.2 n=81, 1.2-1.4 n=167, 1.4-1.6 n=173. Canoe shRNA 1.0-1.2 n=36, 1.2-1.4 n=102, 1.4-1.6 n=116.

## Supplementary Movie 1

**Interphase Long Axis Does Not Predict Division Orientation**

Cells marked with Basigin::YFP to reveal cell outlines and Tubulin-RFP and Centrosomin-RFP to reveal the mitotic spindle and poles. Each frame is a merge of two planes spaced 0.5μM apart. Frames taken one minute apart. The frame rate is seven per second. Scale bar = 5μm.

## Supplementary Movie 2

**Spindle Rotation in a Dividing Follicle Cell**

Tubulin-RFP and Centrosomin-GFP were used to reveal the mitotic spindle and poles. Each frame is a merge of two planes spaced 0.5μM apart. Frames taken one minute apart. The frame rate is seven per second. Scale bar = 5μM.

## Supplementary Movie 3

**Spindle Rotation in the Embryonic Neurectoderm**

Tubulin-RFP and Centrosomin-GFP were used to reveal the mitotic spindle and poles. Each frame is a merge of two planes spaced 0.5μM apart. Frames taken 15 seconds apart. The frame rate is seven per second. Scale bar = 5μM.

